# Mechanical ventilation affects the microecology of the rat respiratory tract

**DOI:** 10.1101/2021.12.07.471701

**Authors:** Chen Xue-Meng, Liu Gao-Wang, Ling Xiao-Mei, Zeng Fan-Fang, Xiao Jin-Fang

## Abstract

**Background:** The most common ‘second strike’ in mechanically ventilated patients is a pulmonary infection caused by the ease with which bacteria can invade and colonize the lungs due to mechanical ventilation. At the same time, metastasis of lower airway microbiota may have significant implications in the development of intubation mechanical ventilation lung inflammation. Thus, we establish a rat model of tracheal intubation with mechanical ventilation and explore the effects of mechanical ventilation on lung injury and microbiological changes in rats.

**Methods:** Sprague-Dawley rats were randomized into control, Spontaneously Breathing (1, 3, 6 hours), Mechanical ventilation(1, 3, 6 hours) groups. Lung wet to dry weight ratio (W/D weight ratio) and Lung histopathological injury score were evaluated.16SrDNA sequencing was performed to explore respiratory flora changes.

**Results:** Bacterial diversity was comparable between healthy and intubation mechanical ventilation rats, with time relation. Ordination analyses revealed that samples clustered more dispersing by tracheal intubation and mechanical ventilation. Finally, predicted metagenomes suggested a substantial increase in biofilm formation phenotype during early tracheal intubation and mechanical ventilation.

**Conclusion:** Collectively, these results establish a link between the duration of mechanical ventilation and alterations to the respiratory tract microecology. In future studies, we hope to discover the effectiveness of new immunomodulatory or probiotic bacteria to prevent airway diseases associated with ventilator therapy.

## 1. Introduction

Ventilator mechanical ventilation is widely used to manage general anesthesia during surgery, respiratory maintenance in intensive care, and perioperative treatment of critically ill patients. However, mechanical ventilation can also lead to lung injury or exacerbate existing lung injury, known as ventilator-induced lung injury (VILI)^1,2^. Also, lung inflammation can occur during ventilator therapy. Current studies have found that mechanical ventilation induces upregulation of cytokine expression in a pro-inflammatory state to the body. Thus, patients are more susceptible to “second strikes” (prolonged mechanical ventilation, aspiration, shock, sepsis, pulmonary infections) ^3^. The most common “second strike” in mechanically ventilated patients is pulmonary infections. The effects of the body’s immune system from the own comorbidities and malnutrition in patients receiving mechanical ventilation under anesthesia6 can also increase the morbidity and mortality of respiratory infections^4^. Respiratory microecology is one of the critical factors in the function of the respiratory tract. Studies over the past few years have demonstrated that the lower respiratory tract is not “sterile” and that, in healthy conditions, the lung microbiota is less dense but harbors a remarkable diversity of interacting microbiota. The ribosomal DNA of Actinobacteria, Aspergillus, Bacteroides, and Bacteroides is present in the lungs of healthy individuals ^5–8^. The “steady state” of the lung microbiome during health may be a process of continuous influx and continuous elimination of unfavorable growth conditions. Studies have confirmed that the balance between immigration and elimination during pulmonary disease is disturbed. The pulmonary microbiota is altered, with bacteria exhibiting competitive dominance ^9^, resulting in an imbalance in the host immune system ^7,10–12^.

The imbalance of respiratory flora may lead to local or even systemic bacterial infections, and the microecological regulatory mechanisms are complex. Identification of microbial colonization by alveolar lavage collected primarily from critically ill patients is currently used clinically to guide anti-infection protocols, while few studies have been reported on the relationship between lung injury and inflammatory response to mechanical ventilation and changes in respiratory flora. Therefore, this study investigated the relationship between lung injury and microbiological changes in rats with tracheal intubation mechanical ventilation and contributed to the study of flora regulation of the respiratory tract in lung injury and inflammation.

## 2. Materials and Methods

To investigate the relationship between lung injury from mechanical ventilation and changes in respiratory flora, we assessed the inflammatory response and microbial changes in the rat airways by pathophysiology and 16SrDNA sequencing.

### 2.2 Animals

Male Sprague - Dawley rats (weighing between 230-330g and aged 8-9 wk) were housed at the Southern Hospital Experimental Centre of Southern Medical University, under the same temperature and humidity environment, and received the same food and water. Animal care and experimental protocols were guidelines by the National Science Council of the Republic of China(NSC1997). The animal studies were reviewed and approved by the Ethical Committee on Animal Experimentation of Nanfang Hospital, Southern Medical University, Guangzhou, China(NFYY-2021-0245).

Twenty-eight male SD rats were randomly divided into seven groups (n=4): control group (group C), tracheal intubation with spontaneous breathing for one hour (group SV1), tracheal intubation with spontaneous breathing for three hours (group SV3), tracheal intubation with spontaneous breathing for six hours (group SV6), tracheal intubation with mechanical ventilation for one hour (group MV1), tracheal intubation with mechanical ventilation for three hours (group MV3), tracheal intubation with mechanical ventilation for six hours (group MV6) (MV6 group). After weighing, we anesthetized the rats with 2% pentobarbital sodium (60 mg/kg) by intraperitoneal injection. We placed our homemade tracheal into the trachea and connected a small animal ventilator (SuperV1.0, HYB, China) for mechanical ventilation. A room temperature of 26-28°C was maintained throughout the procedure. The control group was directly executed after tracheal intubation, while the SV group was intubated and left to breathe independently. The MV group was mechanically ventilated at 1, 3, and 6 hours, respectively. Mechanical ventilation parameters: tidal volume (VT) 7 ml/kg, PEEP = 0 mmHg, respiratory rate 80 breaths/min (adjusted according to respiration during anaesthesia), inhalation-expiration ratio (I: E) = 1:2, inhaled gas was air (oxygen concentration 21%).

### 2.3 Experiments

#### 2.3.1 Rat lung injury assay

##### 2.3.1.1 Lung tissue W/D

To investigate the effect of the duration of mechanical ventilation on the edema of lung tissue, we calculated the wet to dry weight ratio (W/D) value of lung tissue. The tissue was weighed on an electronic balance and recorded as wet weight. The tissue was then baked in an oven at 80°C and weighed to a constant weight, and recorded as dry weight, and the lung tissue was baked for ≥48 h.

##### 2.3.1.2 Histopathological examination of the lung

To observe the pathological damage of lung tissue at different ventilation duration, we performed HE-stained light microscopy of lung tissue. The anterior lobe of the right lung of rats was fixed in 4% paraformaldehyde solution. After paraffin embedding, sectioning, and HE staining, the lung was placed under a light microscope to observe the histopathological changes. Five high magnification views were randomly selected to observe the lung tissue morphology, and the pictures were taken and saved. Scores included lung tissue congestion, septal edema, erythrocyte infiltration, alveolar cavity destruction, capillary destruction, and other pathological changes, and the mean values were taken after scoring ^13,14^.

#### 2.2.4 Detection of respiratory flora in rats

##### Specimen collection

The trachea and left lung of 28 SD rats were removed, snap-frozen in liquid nitrogen, and stored in a -80°C refrigerator to sequence the 16SrDNA colonies.

##### PCR amplification

The 16S rDNA target region of the ribosomal RNA gene were amplified by PCR(95 °C for 5 min, followed by 30 cycles at 95 °C for 1 min, 60 °C for 1 min, and 72 °C for 1 min and a final extension at 72 °C for 7 min)using primers listed in the table ^15^. PCR reactions were performed in triplicate 50 μL mixture containing 10 μL of 5 × Q5@ Reaction Buffer, 10 μL of 5 × Q5@ High GC Enhancer, 1.5 μL of 2.5 mM dNTPs, 1.5 μL of each primer (10 μM), 0.2 μL of Q5@ High-Fidelity DNA Polymerase, and 50 ng of template DNA. Related PCR reagents were from New England Biolabs, USA.

##### DNA sequencing and analysis

Illumina Novaseq 6000 sequencing Amplicons were extracted from 2% agarose gels and purified using the AxyPrep DNA Gel Extraction Kit (Axygen Biosciences, Union City, CA, U.S.) according to the manufacturer’s instructions and quantified using ABI StepOnePlus Real-Time PCR System (Life Technologies, Foster City, USA). Purified amplicons were pooled in equimolar and paired-end sequenced (PE250) on an Illumina platform according to the standard protocols. The raw reads were deposited into the NCBI Sequence Read Archive (SRA) database (Accession Number: SRP******).

#### 2.2.5 Statistical Methods

For the ratio of Lung tissue W/D and lung injury score, One-way ANOVA test was completed. For Observe species index and Simpson index, comparisons were performed by the Turkey HSD test. The significant difference of Beta diversity was tested with Adonis non-parametric test. The Turkey HSD test compared the relative abundance of respiratory microbiota between groups. P-values less than 0.05 were considered statistically significant. One-way ANOVA test was performed by SPSS software, version 25.0 (IBM Corp, New York, NY, USA). The Adonis non-parametric test was performed, and the Turkey HSD test was performed by the OmicShare tools, a free online platform for data analysis (http://www.omicshare.com/tools).

## 3. Results

### Comparison of lung tissue W/D values between groups

The W/D values of lung tissue in the SV and MV groups were significantly higher after three hours. (P< 0.05) (Table 1).

**Table 1.**
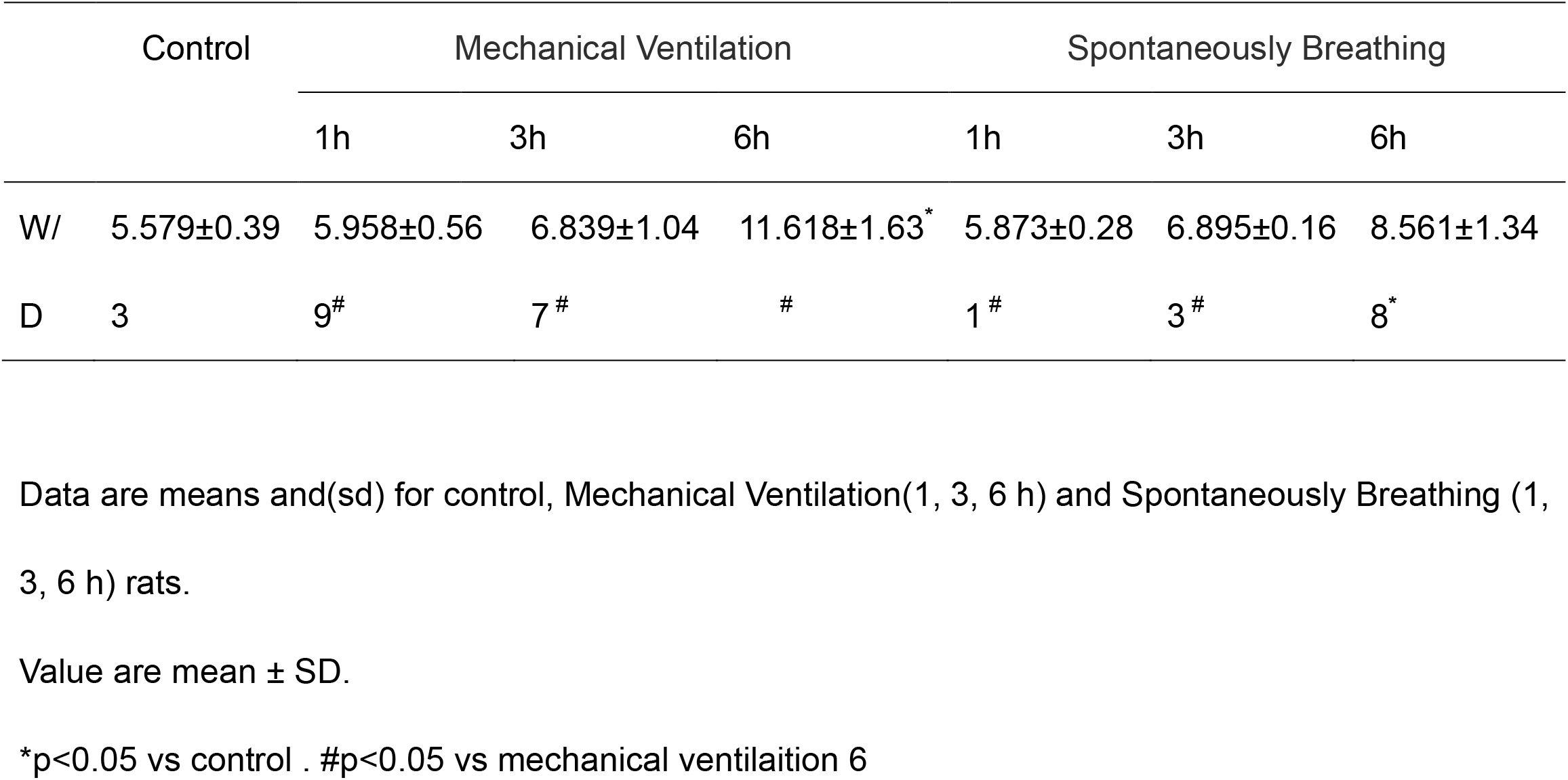
Effect of mechanical ventilation on lung wet to dry weight ratio (W/D) from rats (n=4, 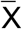±SD)

### Pathological changes in lung tissue

The alveolar structure of group C was normal, with a small number of inflammatory cells infiltrating the lung. Six hours into the SV group, there was significant hemorrhage, widening of the alveolar septum, lymphocytic infiltration, and massive destruction of the alveolar structure. Three hours into the MV group, diffuse intra-pulmonary hemorrhage was seen. Six hours into ventilation, massive diffuse hemorrhage was seen, widening of the alveolar septum, more exudate in the alveolar cavity, destruction of the tissue structure, fusion of the alveoli and solid lung tissue. The degree of lung tissue damage in the SV and MV groups increased with time (Figure 1).

**Figure 1.**
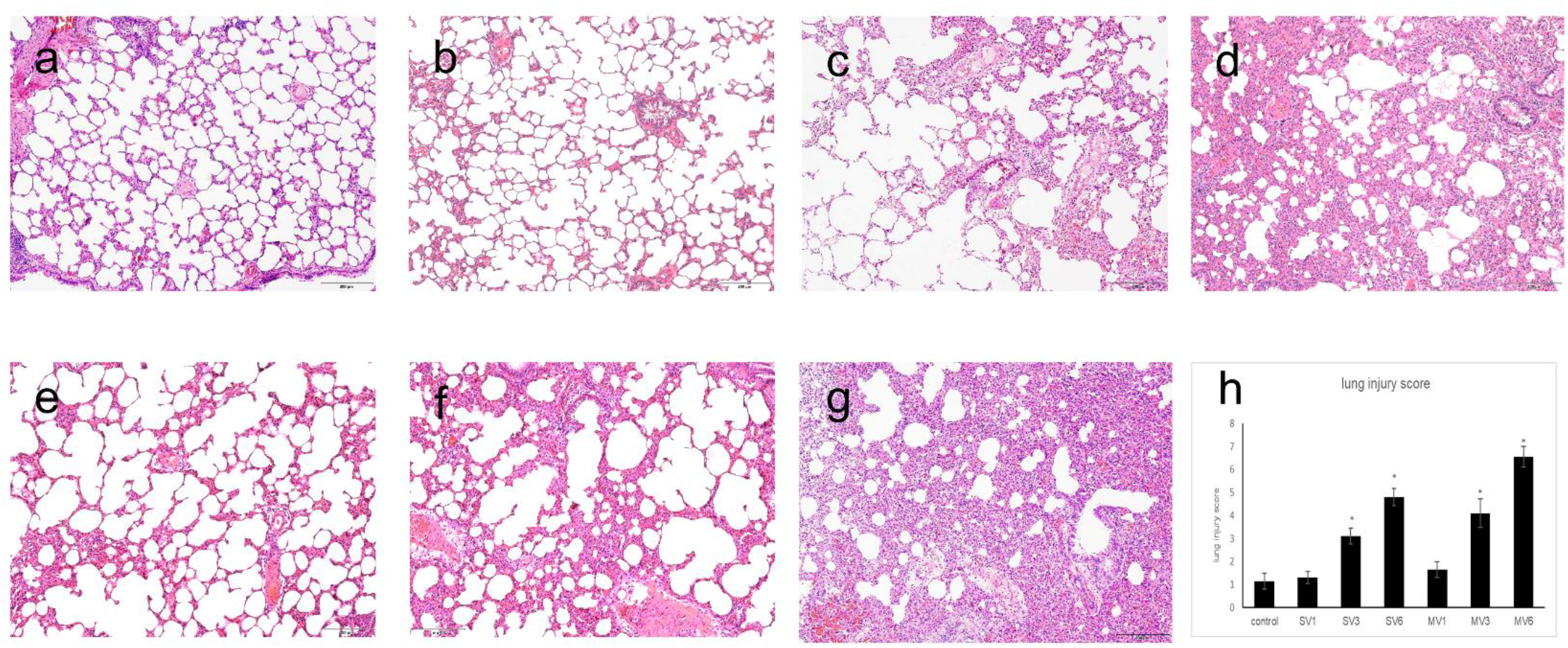
Effect of mechanical ventilation on lung tissues injury from rats. (a) Rats were executed directly after tracheal intubation under anesthesia. (b-d) Rats were kept under anesthesia after tracheal intubation with spontaneous breathing for 1, 3, and 6 h. (e-g). Rats were mechanically ventilated for 1, 3, and 6 h after tracheal intubation. (h) Lung tissue injury score. Data are representative of at least four different experiments. Results are mean ± S,for four rats. *p<0.05, **p<0.01, ***p<0.001, compared with control animals or between mechanical ventilation six hours and spontaneous breathing six hours.

### Analysis of respiratory microbiological changes

#### Analysis of the composition of the respiratory flora

Dominant species largely determine the ecological and functional structure of the microbial community and understand the community’s species composition. We analyzed the bacterial composition using 16S rDNA gene sequencing. After filtering and screening at uniform depth, a total of 24 phyla and 244 genera were identified in the entire sample. To investigate the phylum and genus that accounted for most sequences, we selected species ranked top10 in abundance mean among all samples to show in detail. Others refer to other known species in the figure, and unclassified refer to unknown species ^16,17^. The dominant phyla of rats were Proteobacteria, Firmicutes, Bacteroidetes, Cyanobacteria, Actinobacteria. In healthy rats, the dominant genus was Lactobacillus. In mechanically ventilated rats, the doaminant genus was Acinetobacter at the beginning. The proportion of the dominant genus Acinetobacter decreased, and that of Prevotella_9 increased as the duration of ventilation increased. The predominant genus in the respiratory tract of the rats was also Acinetobacter at the early stage of tracheal intubation, while the proportion of Streptococcus gradually increased with time (Figure 2a. Figure 2b). To further investigate the main flora specific to each group, we applied LEFse to analyze the differences in flora between the seven rat groups ^18^. As shown in the figure(Figure 2c), only the normal rats and the 6hours group showed the main differential flora, Parvibacter for 6 hours of autonomic breathing, Megamonas for 6hours of mechanical ventilation, and normal rats Acinetobacter_radioresistens.

**Figure 2.**
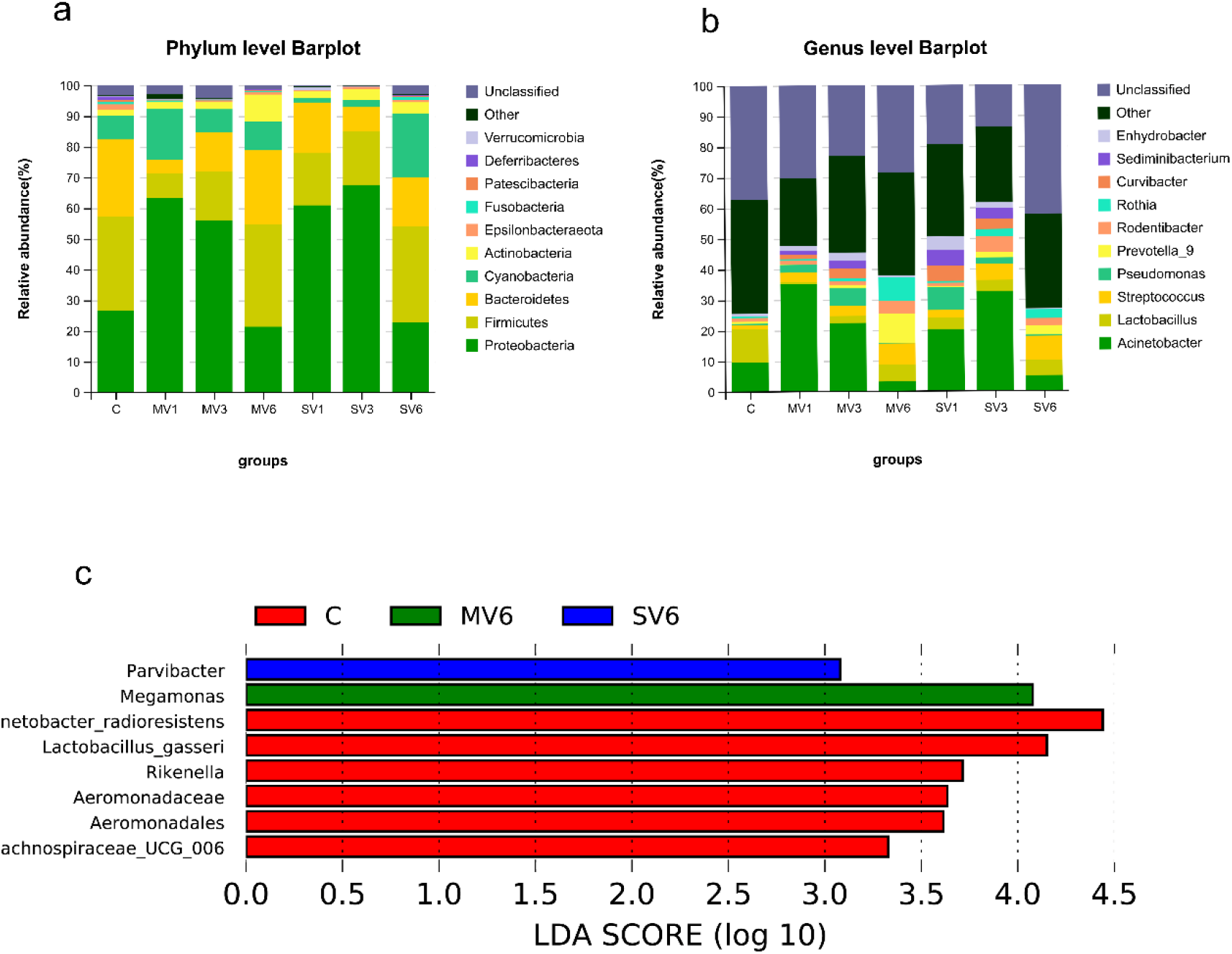
Changes in the lower respiratory tract flora of rats during mechanical ventilation. (a) At the phylum level, the main dominant phyla were Proteobacteria, Firmicutes, Bacteroidetes, Cyanobacteria, and Actinobacteria. (b) At the genus level, the main dominant genera were Acinetobacter, Lactobacillus, and Staphylococcus. Streptococcus. (c) LEFse analysis (LDA>2.0) was performed for all groups, with corresponding differential strains for control, MV6, SV6, Acinetobacter_radioresistens for control, Megamonas for MV6, and Parvibacter for SV6. In summary, tracheal intubation mechanical ventilation caused changes in the lower respiratory tract flora ratio in rats.

As expected, the two operations of tracheal intubation and mechanical ventilation resulted in alterations in the respiratory microecology of rats. The main strains of normal adult BALF are Actinobacteria, Proteobacteria, Bacteroidetes, and Firmicutes ^5–8^. The strains of rats in this study cover the main human strains and are similar to humans. It has been demonstrated that representatives of the genus Acinetobacter come from the lung microbiome of mechanically ventilated patients with suspected pneumonia ^19^. In the present study, the elevated proportion of Acinetobacter as the dominant species during the initial phase of mechanical ventilation with tracheal intubation may have contributed to the gradual pathological damage of lung tissue. As the duration of ventilation increased, the proportion of Prevotella_9 gradually increased, and it was considered that the subclinical lung inflammatory manifestations caused by mechanical ventilation created an environment for Prevotella_9 to be retained in the lungs^20,21^. Two classes of probiotics, Parvibacter and Megamonas, are often present in the intestinal tract of humans and animals, and the induction of mechanical ventilation by tracheal intubation itself may be associated with changes in intestinal microbial composition and diversity. There is now strong evidence that the gut microbiota can influence the microbiology of the lung through direct inoculation of bacteria or the distribution of SCFAs ^22,23^. Characterization of lower airway probiotic bacteria associated with lung injury from tracheal intubation with mechanical ventilation may be necessary for in-depth exploration of methods to prevent such injury.

Interestingly, although many taxa are shared between samples, the absence of pathogens (e.g., dominant genera) in one sample does not predict their absence in paired sample sequence data. For example, samples V1-1 and V3-1, despite not detecting Sediminibacterium and Curvibacter, showed trends consistent with other genus compared to controls. It is important to note that the dominant genus in samples V1-2 was Streptococcus and the second-highest proportion was Acinetobacter. These examples show that different pathogens can vary in different individuals of the same organism, and therefore, 16S rDNA sequence data are not always representative of lower respiratory tract infections. Lower airway Bacterial diversity varies in healthy and ventilator-assisted breathing rats.

#### Alpha Diversity

As mentioned above, in each sample, a few taxa dominate the majority of the sequences. Two alpha diversity metrics, Observed species, and Simpson diversity were used to investigate this issue further to measure species richness and homogeneity ^24,25^. The OTU was highest in the mechanically ventilated 6hours group, indicating the highest abundance. However, the difference between the control and autonomic breathing 6hours groups was not significant, and OUT abundance decreased at 1hour and 3hours in both ventilated and autonomic breathing groups (Figure 3a). Using the Simpson diversity index, the abundance and homogeneity of the respiratory flora decreased in each experimental group compared with the control group, but the differences were not significant (Figure 3b and Figure 3c). These data suggest that mechanical ventilation tracheal intubation alters the abundance and homogeneity of lower airway microecology with time influenced.

**Figure 3.**
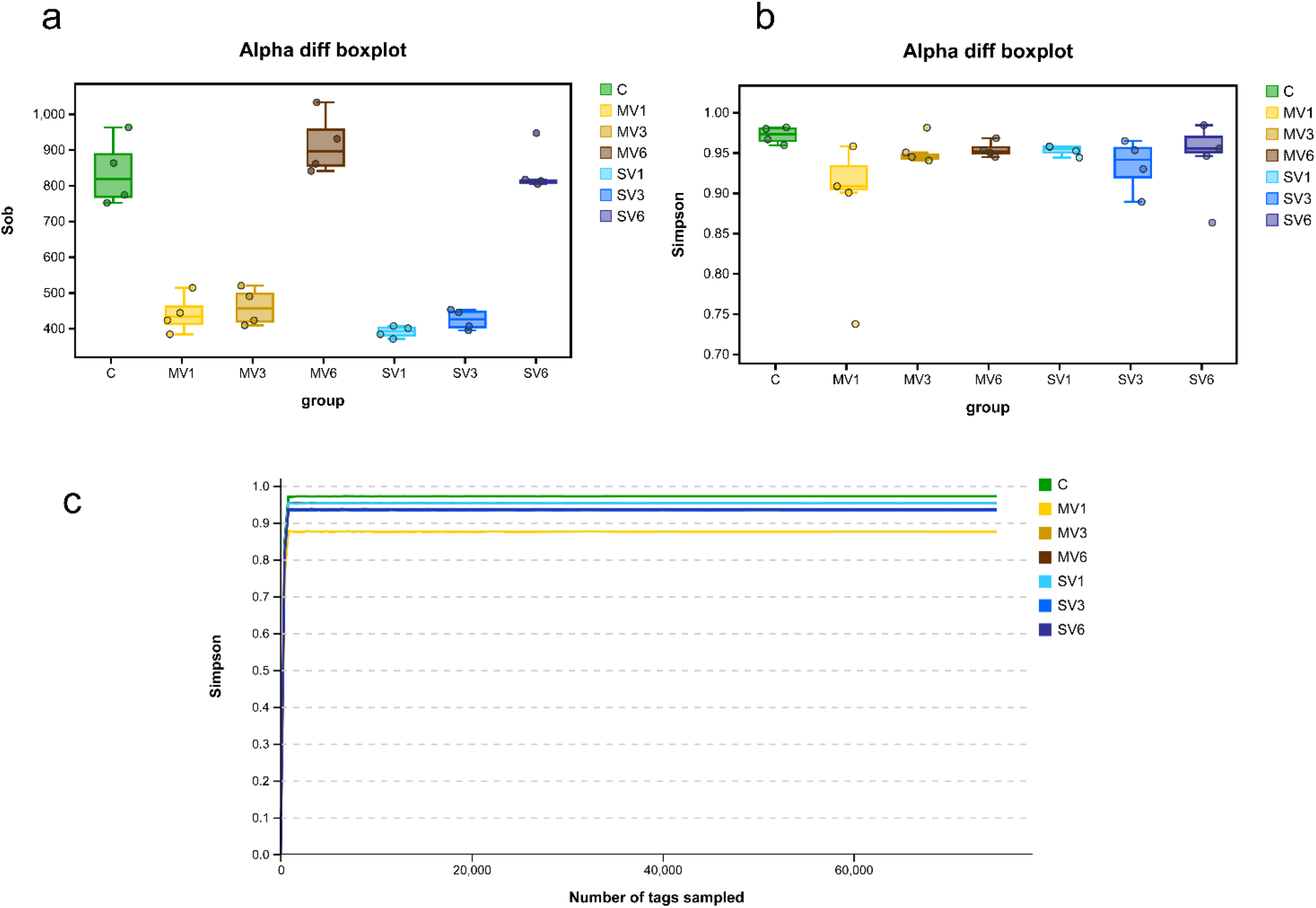
Alpha diversity of rat lower respiratory tract flora. (a) The observed species index decreased in the first 3 hours of mechanical ventilation and gradually increased as ventilation was prolonged. (b) Simpson index was not statistically different among groups, and the homogeneity of species was more consistent among groups. (c) Alpha diversity dilution curves leveled off in all groups, indicating a sufficient amount of data in each group.

#### Beta Diversity

Beta diversity of the respiratory microbiota was visualized using a ranking method. We were interested in whether the bacterial community aggregates tightly due to tracheal intubation or mechanical ventilation. Previous evidence suggests that three main factors determine the composition of the lower respiratory microbiota, namely microbial migration, microbial elimination, and the relative replication rate of its members.^9^. The oral and pulmonary have similar microbiomes community^1,9,26^. A study has investigated that the oral microbiome is part of the homeostatic processes regulating lung inflammation ^26^. Mechanical ventilation operations with tracheal intubation may dysregulate microecology by impairing lower airway defense mechanisms ^27,28^. We hypothesized that samples are more intensively clustered by tracheal intubation and mechanical ventilation based on previous studies. To confirm this hypothesis, we used unweighted Unifrac and weighted Unifrac, two abundance-sensitive, phylogenetically relevant diversity metrics, to compare samples^29^. The two metrics were calculated to determine the pairwise phylogenetic distances between each sample, then plotted using principal coordinate analysis (PCoA) ^30^. From the unweighted Unifrac, contrary to our hypothesis, samples from the SV and MV groups did not cluster more closely with healthy rats (PERMANOVA, P=0.001) (Figure 4a). The graph shows that the mechanically ventilated group (PERMANOVA, R2=0.3737, P=0.002) and the autonomously ventilated group (PERMANOVA R2=0.4702, P=0.002) both exhibited moderate phylogenetic differences(Figure 4b,Figure 4c). Although experimental groups were better aggregated with healthy rats in terms of weighted Unifrac (PERMANOVA, P=0.122), the autonomic breathing group exhibited more significant phylogenetic differences than the mechanically ventilated group (PERMANOVA, R2=0.6939, P=0.001) (Figure 4d, Figure 4e, Figure 4f). These analyses suggest that the overall community structure of the rat lower airway is influenced by tracheal intubation and mechanical ventilation.

**Figure 4.**
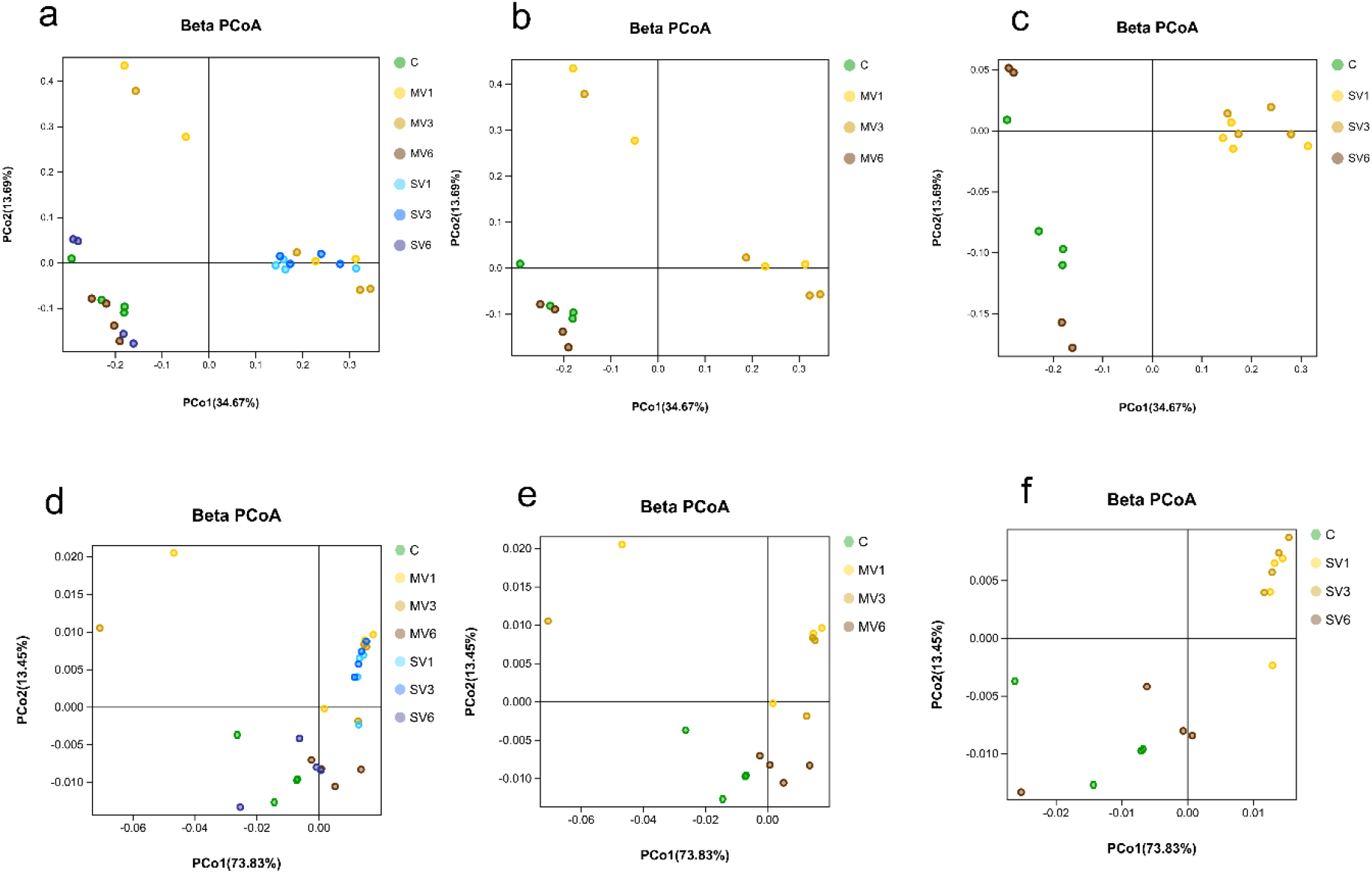
Beta diversity of rat lower respiratory tract flora. (a-c) In unweighted Unifrac analysis, tracheal intubation mechanical ventilation made the rat’s lower respiratory flora more fragmented. (d-f) In the weighted Unifrac analysis of all groups, tracheal intubation mechanical ventilation did not make the rat respiratory flora more dispersed. However, in comparing normal rats with the spontaneously breathing group, a more significant dispersion of the rat lower respiratory flora occurred.

#### Predicted functional categories(功能预测分析)

To gain insight into the function of the bacterial community, PICRUSt2^31^ was implemented to infer bacterial metagenomes based on 16S rDNA gene content. There were 33 unique KEGG pathways among the inferred metagenomes and showed more remarkable similarities between normal rats and each ventilated group (Figure 5). This shows that mechanical ventilation by tracheal intubation does not significantly alter the functional capacity of the bacterial community in the lower respiratory tract of rats. To further summarize the output of PICRUSt, we utilized BugBase, a bioinformatics tool that allows inference of community-wide phenotypes from predicted metagenomes^32^.BugBase found that gene function associated with anaerobiosis was enriched in the lower airway samples and, interestingly, gene function was reduced early in mechanical ventilation (CvsMV1 P= 0.013). The important genus attributed changed, with Proteobacteria being the dominant genus for 1hour of mechanical ventilation, and this dominance tended to decrease with time (Figure 6A). Biofilm formation and Gram-negative phenotypes were bacterial genus phenotypes frequently associated with respiratory pathogenicity. Our study suggested that biofilm formation phenotypes were substantially elevated during early tracheal intubation and mechanical ventilation, with Proteobacteria being the predominant cause of this phenotype. Moreover, Gram-negative phenotypes have no significant differences between groups, but phenotypes were more diverse in samples from 6hours of mechanical ventilation. Biofilm-forming bacteria were observed in the predicted metagenomes of each group. With the increasing understanding of the microbiota, we recognize interactions between the microbiota and the immune system of the airway epithelium. For example, asthma and allergy^11^ vary with the occurrence of airway infections. Studies have confirmed that microinhalation, external pressure, etc., favor the establishment of microbial biofilms. Biofilms have significant antimicrobial tolerance, evading host immune system attack and persisting in the host ^33^. In the present study, biofilm-associated gene phenotypes were enhanced in rats undergoing mechanical ventilation with tracheal intubation, indicating enhanced biogenesis, consistent with previous studies ^34^. Overall, these data suggest that the initial phase of mechanical ventilation by tracheal intubation disrupts the ecotone. Anaerobic phenotypic function decreased, and other phenotypes associated with bacterial pathogenicities also were influenced. However, it is also difficult to explain whether the reduced diversity of the microbiome is a causative factor for disease or whether diversity is simply a response to disease inflammation.

**Figure 5.**
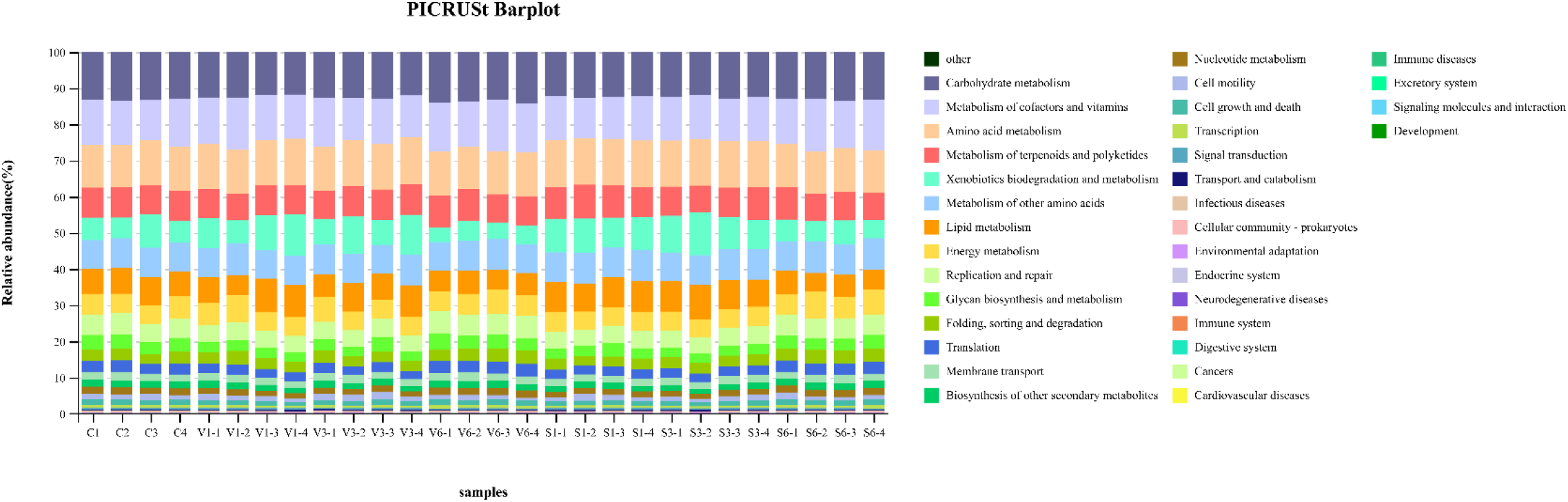
PICRUSt2 analysis of rat lower respiratory tract flora. Thirty-three unique KEGG pathways among the inferred metagenomes. It is showed more remarkable similarities between normal rats and each ventilated group.

**Figure 6.**
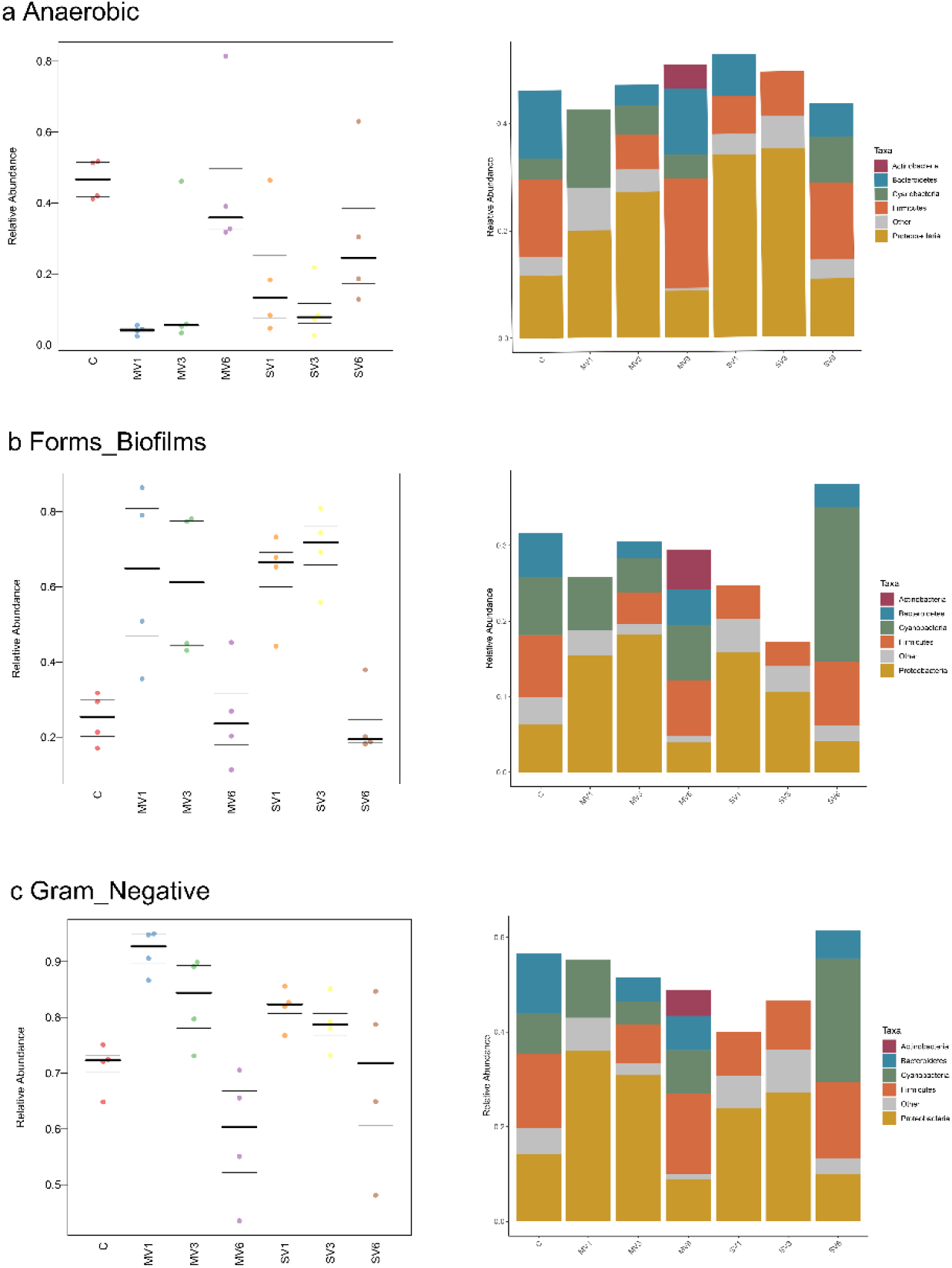
Bugbase analysis of rat lower respiratory tract flora. (a) With the associated representative genera changes, anaerobic activity is diminished at the beginning of mechanical ventilation by tracheal intubation. (b) Forms_Biofilm is enhanced at the beginning of mechanical ventilation by tracheal intubation, and the associated representative genera are altered. (c) Gram_Negative formation did not vary much between groups, slightly increased at the beginning of the experiment, and the proportion slightly decreased with the prolongation of the experiment.

## 4. Discussion

With the continuous medical advances, tracheal intubation mechanical ventilation has been widely used in managing general anesthesia during surgery, intensive care units, and emergency resuscitation of critically ill patients. However, mechanical ventilation is also a “double-edged sword,” which may lead to lung injury or aggravate existing lung injury ^35,36^. Volume injury due to lung hyperinflation and shear injury due to repeated opening and closing of the terminal airway play an essential role in the pathogenesis of VILI^3^. On the other hand, respiratory inflammation may occur during ventilatory therapy. As an open cavity between the human body and the outside world, the respiratory tract has a normal microbiota that is an integral part of the human body’s natural defense. Respiratory microbiota provides a biological barrier against the invasion of foreign bodies or pathogenic bacteria through mechanisms such as spatial occupancy, healthy competition, and secretion of antibacterial or bactericidal substances^37–40^. As one of the most frequently contacted parts of the human body, the respiratory micro-ecosystem is essential in maintaining human health and is also most vulnerable to damage. In recent years, the lower respiratory tract microbiota has been extensively studied and reported to be similar to that of the oropharynx^1,2,41^. However, studies exploring the relationship between mechanical ventilation and lower airway microecology have mainly explored at the level of ventilator-associated pneumonia (VAP) ^42–45^. Changes in lower airway microecology during the initial phase of mechanical ventilation have not been reported. To investigate the in vivo response of tracheal intubation mechanical ventilation on lower respiratory microecology in SD rats in this study, we obtained pathological findings of lung inflammation and injury and data on bacterial community structure.

In the present study, we observed that three hours after the administration of mechanical ventilation with tracheal intubation, hemorrhagic spots appeared in the lung tissue, widening of the alveolar septum, large amounts of exudate in the alveolar cavity, and altered alveolar structural damage, which aggravated with time. Correspondingly, the highest percentage of Proteobacteria was present in the experimental and control groups. At the onset of mechanical ventilation, the percentage of Proteobacteria in the lower airways increased more than twofold compared to controls. At the same time, the number of Firmicutes and Bacteroidetes decreased about fourfold. When six hours of mechanical ventilation had elapsed, Actinobacteria was more than four times that of the control group. These results suggest that mechanical ventilation with tracheal intubation alters the structure of the microecology while inducing lung tissue damage. However, how bacterial changes and tissue damage interact and require further study. We observed a significant amount of Acinetobacter within three hours of mechanical ventilation; therefore, Acinetobacter may be one of the factors involved in mediating the proinflammatory phase of lung tissue injury. It has been demonstrated that Acinetobacter may be the predominant bacteria in the lung microbiota of mechanically ventilated patients with pneumonia^19^. The excessive growth of this pathogen may promote airway injury and exacerbate microecological dysregulation. Lactobacillus and Streptococcus gradually increase with prolonged ventilation. Lactobacillus and Streptococcus act as probiotic strains. On the one hand, altering the composition of the host-microbiome dysregulate the microecology. On the other hand, they act indirectly through the standard mucosal immune system (CMIS) ^46^. These probiotic strains may play a positive role in alleviating respiratory impairment and the corresponding microecological imbalance. In the mechanically ventilated 6-hour group, the main differential bacteria genus Prevotella_9 likewise belongs to probiotic bacteria and is often present in the intestine. Current studies have confirmed the existence of an intestinal-pulmonary microbiome axis in the organism^47^. AS the metabolically active bacteria, Bacteroides and Firmicutes produce SCFA. We know that the gut microbiota can influence the microbiology of the lung through the distribution of SCFAs. SCFAs play a key role in maintaining mucosal immunity in the gut and lung tissues as a bridge between the microbiota and the immune system. ^22,23^. Our findings may be relevant to understanding the mechanisms between tracheal intubation mechanical ventilation and lung tissue injury. Mechanical ventilation by tracheal intubation leads to an inflammatory response characterized by impaired airway defense mechanisms and alters in the microenvironment structure of the lower airways. In the future, we will further investigate probiotics to assess the prognostic relevance of microbial imbalance and the effectiveness of specific therapeutic interventions.

We analyzed lower airway microorganisms’ alpha and beta diversity. We found that both tracheal intubation and mechanical ventilation cause changes in the microbial composition of the lower airways. At the beginning of tracheal intubation mechanical ventilation, bacterial species diversity decreased, indicating that intubation ventilation impaired the ecological niche balance in the lower airways of rats and disturbed bacterial diversity. Six hours after ventilation, bacterial diversity was elevated, and species composition remained different from healthy rats, considering that it may be related to pathogen overgrowth and stimulation of the host immune system that promotes disease development^48^. It is noteworthy that the experimental and control groups differed in bacterial species abundance but were more homogeneous. As seen in this study, the samples from the experimental and control groups were not well aggregated, suggesting that the overall community structure of the rat lower airway is influenced by tracheal intubation and mechanical ventilation. However, the sample size of the groups in this experiment was small, and there were subject differences between individuals^37,39,49^. The sample size needs to be increased in subsequent experiments to explore the relationship between the two further.

There may be a relationship between mechanical ventilation status and airway immune function. Previous studies have shown that lower respiratory microbes are essential in the immune system’s maturation, education, and function. Especially, the regulation of the inflammatory response in respiratory diseases such as asthma^50^, chronic obstructive pulmonary disease (COPD) ^51^, cystic fibrosis^52^, and respiratory infections^53^ plays a key role. Ecological dysregulation leads to overgrowth of pathogenic bacteria, loss of symbiotic microbial diversity, and, ultimately, an inflammatory response in the host, leading to disease ^54 55^. In the present study, after stimulation by mechanical ventilation with tracheal intubation, the microecological balance in the lower airways was disturbed, the loss of symbiotic microbial diversity, the immune system was affected, and inflammatory pathological changes began to develop in the lung tissue. The inflammatory cascade response and pro-inflammatory cytokines are hyperactivated with prolonged mechanical ventilation. Eventually, lung tissue increased damage. In BugBase functional analysis, we found biofilm formation gene-function phenotypes elevated abundance compared to the control group. It may be associated with damage to airway epithelial cells and disruption of defense mechanisms due to stimulation by tracheal intubation mechanical ventilation maneuvers. Biofilms have significant antimicrobial tolerance and can persist in the host and evade the host immune system ^33,56^. We know that specialized respiratory epithelial cells are necessary to maintain immune homeostasis in the lower airway tract. As the first line of defense against potentially harmful environmental stimuli, respiratory epithelial cells are involved in maintaining immune homeostasis. Thus, epithelial cell dysfunction may influence the development of many airways and pulmonary inflammatory diseases^57,58^. In addition, it has been demonstrated that NF-κB, TINCR, and PGLYRP4 (rs3006458) are involved in airway immune mechanisms^59,60^. In future studies, we hope to investigate further the relationship between the mechanical ventilation status of tracheal intubation alone and epithelial cell alterations. We will make predictions of relevant metabolic pathways by metatranscriptomic metabolomic and proteomic studies.

In conclusion, our findings may be relevant to understanding the altered immune system mechanisms between tracheal intubation mechanical ventilation and lung tissue injury. Such operations as tracheal intubation ventilator-assisted ventilation lead to lung injury and inflammatory responses characterized by changes in the microbial community of the lower airways that become temporally correlated. The present study provides new ideas for functional studies on the role of pulmonary commensals in airway morphogenesis development, epithelial homeostasis, and the immune system. Particularly, it allows a more systematic study of bacteria’s colonization and growth characteristics on pulmonary epithelial cells, their impact on lung infection and inflammation. Future studies hope to discover the effectiveness of new immunomodulatory or probiotic bacteria to prevent airway diseases associated with intraoperative or postoperative short-term intubation.

## Acknowledgements

We thank the Animal Experimentation of Nanfang Hospital, Southern Medical University (Guangzhou, China) for performing animals and experimental sites. We are grateful for the sequencing platform or bio information analysis of Gene Denovo Biotechnology Co., Ltd (Guangzhou, China).

